# Atypically larger variability of resource allocation accounts for visual working memory deficits in schizophrenia

**DOI:** 10.1101/424523

**Authors:** Yi-Jie Zhao, Tianye Ma, Xuemei Ran, Li Zhang, Ru-Yuan Zhang, Yixuan Ku

## Abstract

Schizophrenia patients are known to have profound deficits in visual working memory (VWM), and almost all previous studies attribute the deficits to decreased memory capacity. This account, however, ignores the potential contributions of other VWM components (e.g., memory precision). Here, we measure the VWM performance of schizophrenia patients and healthy control subjects on two classical delay-estimation tasks. Moreover, we thoroughly evaluate several established computational models of VWM to compare the performance of the two groups. We find that the model assuming variable precision across items and trials is the best model to explain the performance of both groups. According to the variable-precision model, schizophrenia subjects exhibit abnormally larger variability of allocating memory resources rather than resources per se. These results invite a rethink of the widely accepted decreased-capacity theory and propose a new perspective on the diagnosis and rehabilitation of schizophrenia.

---

Schizophrenia is a severe mental disorder accompanied by a range of dysfunctions in perceptual and cognitive behavior, among which working memory deficit is considered as a core feature ^1–4^. Working memory refers to the ability to temporally store and manipulate information in order to guide appropriate behavior, and it has been shown to link with a broad range of other brain functions, including perception, attention, problem-solving and executive control ^5–8^. Dysfunctions in working memory therefore might cascade into multiple mental processes, causing a wide spectrum of negative consequences ^2,3,9^.

A well-established finding in lab-based experiments is that people with schizophrenia (SZ) exhibit worse performance than healthy control (HC) in visual working memory (VWM) tasks ^2^. This phenomenon has long been attributed to decreased VWM capacity in SZ ^2,10,11^. The theory of decreased capacity was supported by previous studies using various VWM or other WM tasks, including the ‘span’ tasks (e.g., digit span, spatial span, verbal span) ^12,13^, the N-back task ^14–16^, the delayed-response task ^17–19^, the change detection task ^20–24^, and the delay-estimation task ^25–27^. Despite the considerable differences across behavioral tasks, almost all previous studies converged to the same conclusion that decreased capacity is the major cause of the VWM deficits in SZ.

Besides capacity, people have increasingly recognized memory *precision* as another pivotal determinant of VWM performance in the basic research of VWM ^28^. Precision reflects the amount of memory resources assigned to individual items—a larger amount of resources leads to higher memory precision. At the neural level, low perceptual precision might arise from either the intrinsic noise in neural processing ^29–31^ or the fluctuations of cognitive factors (e.g., arousal, attention) ^31,32^. Atypically increased variability in both behavioral and neural responses has been discovered in patients with mental diseases such as autism spectrum disorder ^33,34^, dyslexia ^35^, and attention-deficit/hyperactivity disorder ^36^. These theoretical and empirical studies raise the possibility that SZ and HC might differ in memory precision rather than capacity—that is, these two groups might be able to remember an equal number of items (i.e., comparable capacity) but SZ generally process and maintain items in a less precise manner. Only a few studies have attempted to simultaneously quantify memory capacity and precision in schizophrenic or schizotypy subjects, and the results did not reach a consensus ^25,26^.

Despite the confound of different VWM components as the possible causes, it is unclear whether SZ and HC employ the same computational strategies (i.e., observer model) in VWM. Most prior studies only used one model and implicitly assumed that model is the best one for both SZ and HC. But without systematic model comparisons model optimality cannot be firmly warranted, and endowed results might be biased by the choice of a particular model. Given that several influential models have been proposed to explain the VWM behavior in normal subjects ^28^, it remains unclear which one is the best for SZ. If the best model for SZ differs from the one for HC, it indicates that the two groups use qualitatively different computational strategies to complete behavioral tasks. Otherwise, if SZ and HC share the same best model, it indicates that they use the same memory structure but quantitatively different parameters. These possibilities should be thoroughly tested using transparent model comparisons.

In the present study, we aim to systematically disentangle the impact of memory capacity and precision, as well as other factors (i.e., variability in allocating resources and variability in choice) in SZ. In this study, the performance of SZ and demographically matched HC was measured in two standard VWM delayed-estimation tasks (Fig. 1). To obtain generalizable results, we tested separate groups of subjects on the delay-estimation paradigm using two independent visual features—color and orientation. Using standard tasks allows us to compare our results with those from previous studies ^25,37–40^. Most importantly, in contrast to most prior studies, we evaluated and compared several mainstream computational models in visual working memory research. This approach allows us to take an unbiased perspective and search a large space of both models and parameters. We believe that the well-controlled tasks and thorough computational modeling will shed new light on the mechanisms of VWM deficits associated with schizophrenia.

**Figure 1.**
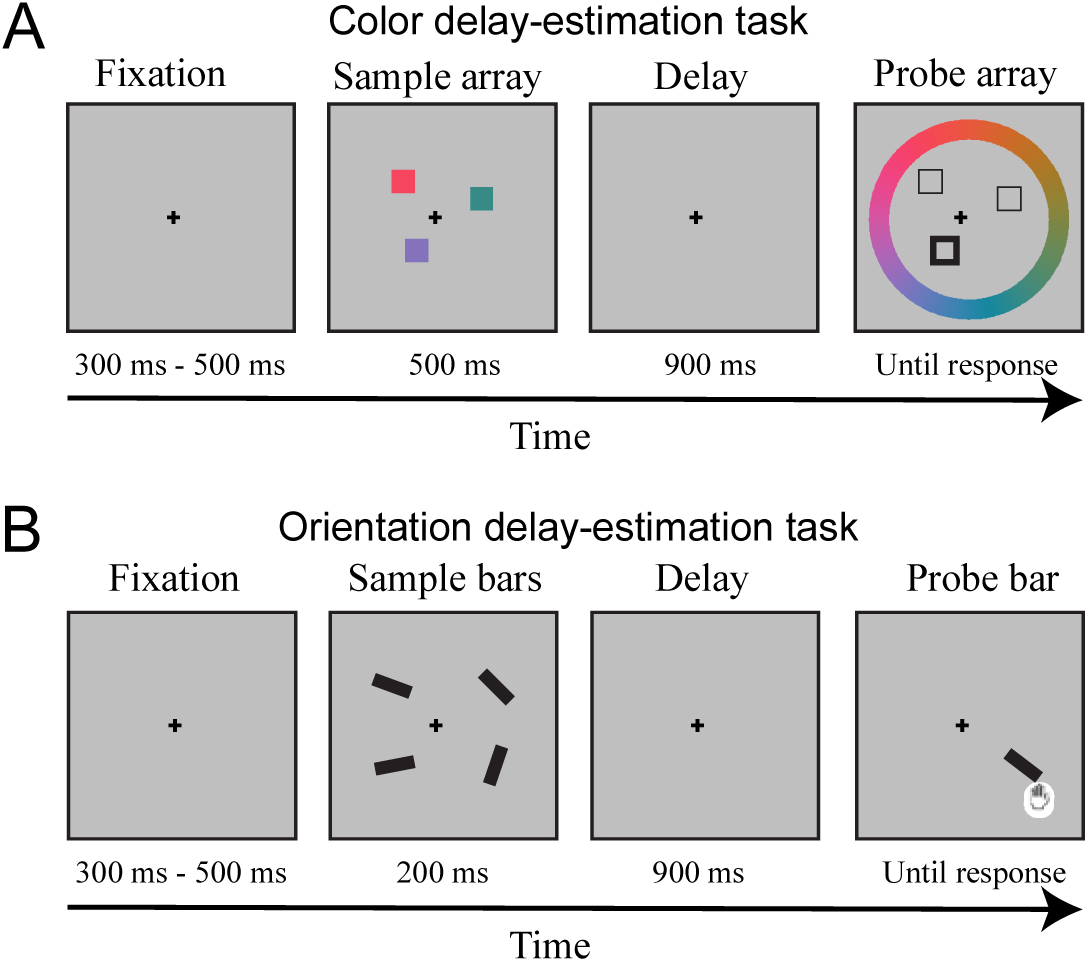
Schematics of the color (***panel A***) and the orientation (***panel B***) delay-estimation experiments. In the color experiment, subjects are instructed to first memorize the colors of all squares (i.e., set size = 3 in this example trial) on the screen, and after a 900ms delay choose the color of the probed square (the one in the left lower visual field in this example) on a color wheel. The orientation experiment is similar to the color experiment. Subjects are instructed to remember the orientations of a set of bars (i.e., set size = 4 in this example trial) and then adjust the orientation of the probed bar using a mouse. Both color and orientation stimuli can be described using a circular feature space of (0, 180]. Response error is the difference between the reported color or orientation and the real color or orientation of the probe in this standard feature space.

## Results

### Worse VWM performance in SZ in both color and orientation delay-estimation experiments

We tested subjects on two delay-estimation experiments. In the color delay-estimation experiment, subjects viewed a set of squares on the screen, and were instructed to memorize their colors. After a short blank, subjects were asked to report the color of one of the squares (i.e., probe) by clicking the color on a color wheal. In order to test a large cohort of subjects, we reduced the difficulty of the task and tested them on two set size levels (i.e., the number of items to memorize). Similarly, in the orientation delay-estimation experiment, subjects were instructed to remember the orientations of a set of bars and then manually adjusted the orientation of the probe bar to the memorized orientation. To improve the efficacy of computational modeling, we increased the task difficulty and tested subjects on set size levels 1, 2, 4, 6 (see experimental details in Methods). These two tasks have been frequently used in prior VWM research ^37,41,42^. Note that we recruited distinct groups of subjects for the two tasks in order to avoid task-or subject-specific confounding factors.

The precision of memory in each trial can be quantified as the difference between the reported color (or orientation) and the true color (respectively, orientation) of the probe. For the color experiment, a repeated-measure ANOVA was performed with circular standard deviation (CSD) as the dependent variable, set size (1/3) as the within-subject variable, group as the between-subject variable (Fig. 2A). Consistent with findings in previous literature, VWM performance was worse when set size was higher (F(1, 119) = 641,703, p < 0.001, partial η^2^ = 0.844), and HC performed significantly better than SZ (F(1,119) = 13.651, p < 0.001, partial η^2^ = 0.103). The interaction between set size and group was not significant (F(1,119) = 0.229, p = 0.633, partial η^2^ = 0.002).

**Figure 2.**
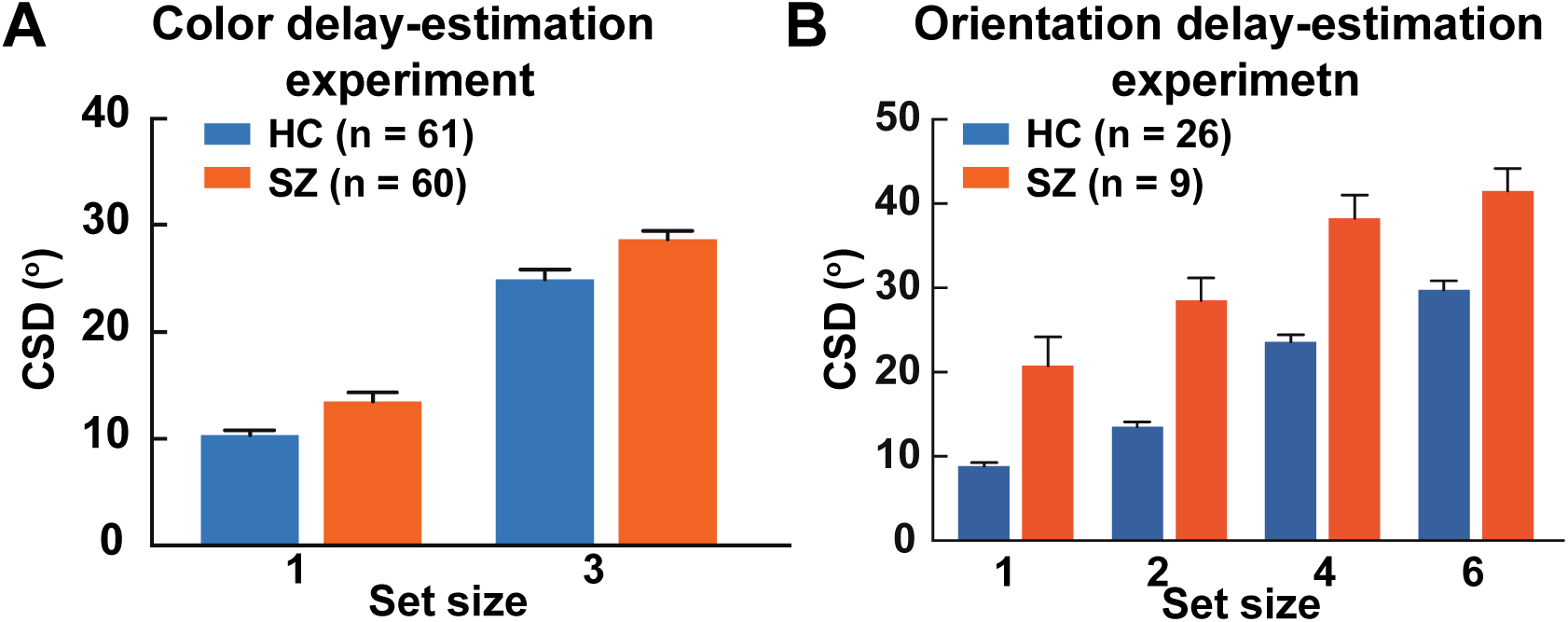
Behavioral performance of SZ and HC in the two tasks. Circular standard deviations of response errors in color (panel A) and orientation (panel B) delay-estimation tasks. SZ show higher CSDs (i.e., worse performance) than HC. All error bars represent SEM across subjects.

In the orientation experiment, similarly, we also found that HC achieved significant lower CSD ((F(3, 31) = 62.967, p < 0.001, partial η^2^ = 0.656). Unsurprisingly, both groups performed worse as set size increased (F(3,31) = 87.682, p < 0.001, partial η^2^ = 0.895). The group by set size interaction was significant (F(3,31) = 3.043, p = 0.043, partial η^2^ = 0.228), with smaller group differences on set size 1 and 6 and slightly larger group differences on set size 2 and 4 (p < 0.001 for all pairwise comparisons, Bonferroni corrected).

Taken together, we replicated the widely documented VWM deficits in SZ in both color and orientation delay-estimation tasks.

### Variable-precision model accounts for VWM behavior in both HC and SZ

To systematically compare the VWM performance between SZ and HZ, we evaluated several mainstream computational models of VWM. We provide some brief introductions here, and readers may consider to skip the following paragraph to directly reach the result part or go to Supplementary Materials for detailed mathematical and intuitive explanations of the models, depending on reading preference.

The first one is the item-limit (IL) model. The IL model assumes no uncertainty in the sensory encoding stage, and that each subject has a fixed memory capacity and a fixed response variability across set size levels ^43^. The second one is the mixture (MIX) model, similar to the IL model but assuming response variability is set-size dependent ^25,26^. Compared with the MIX model, the slots-plus-averaging (SA) model ^37^ further elaborates the idea that memory resources manifest as discrete chunks, and these chunks can be flexibly assigned to multiple items. We also explored a modified version of the SA model, dubbed cosSA model, which inherits the idea of discrete memory resources and further assumes that response bias is stimulus-dependent and can be described as empirically derived periodic functions. The fifth one is the equal-precision (EP) model, which is similar to the variable-precision (VP) model (Fig. 4) below but assumes that the memory resources are evenly distributed across items and trials ^44,45^. The VP model proposes that memory resources are continuous, and the amount of resource assigned to individual items varies across items and trials. Note that the VP model does not include the capacity component thus we also constructed a variable-precision-with-capacity (VPcap) model that not only acknowledges the variable precision mechanisms but also explicitly estimates capacity of individual subjects. Note that the IL, MIX, SA and cosSA, and VPcap models have the parameter of capacity, and the EP and VP models do not. Here, capacity is operationally defined as the maximum number of items that can be stored in memory. Some items are out of memory if set size exceeds capacity, and the subject has to randomly guess the color if one of the out-of-memory items is probed.

**Figure 3.**
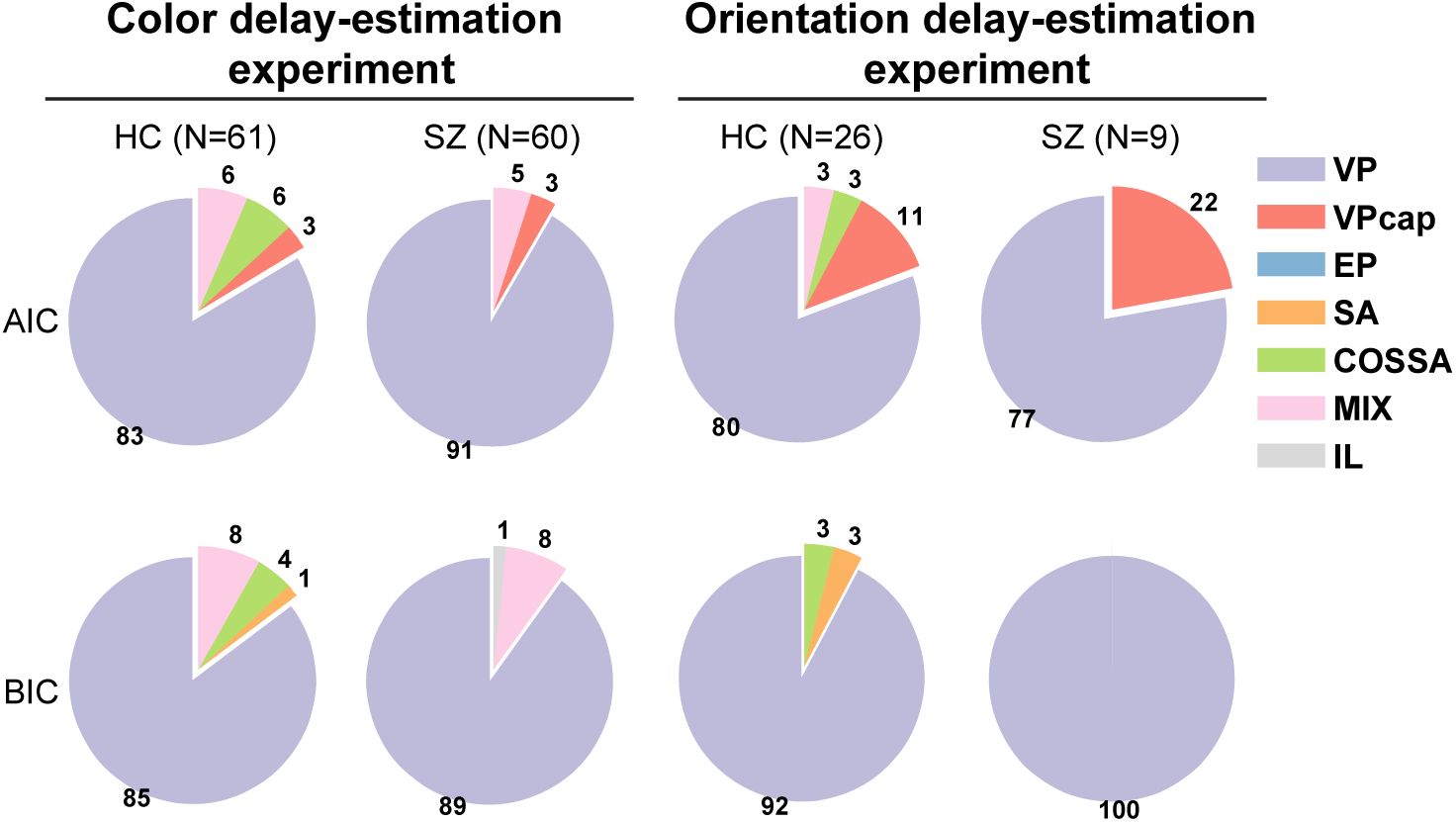
Model comparison results. A total of seven computational models are compared in each subject. The pie charts illustrate the proportion (i.e., the percent number shown in each slice) of subjects for whom each model is their best-fitting model. Under both AIC and BIC model comparison metrics, the VP model is the best-fitting model for the majority of subjects in both groups and in both experiments. This result indicates both groups share a qualitatively similar internal process of VWM.

**Figure 4.**
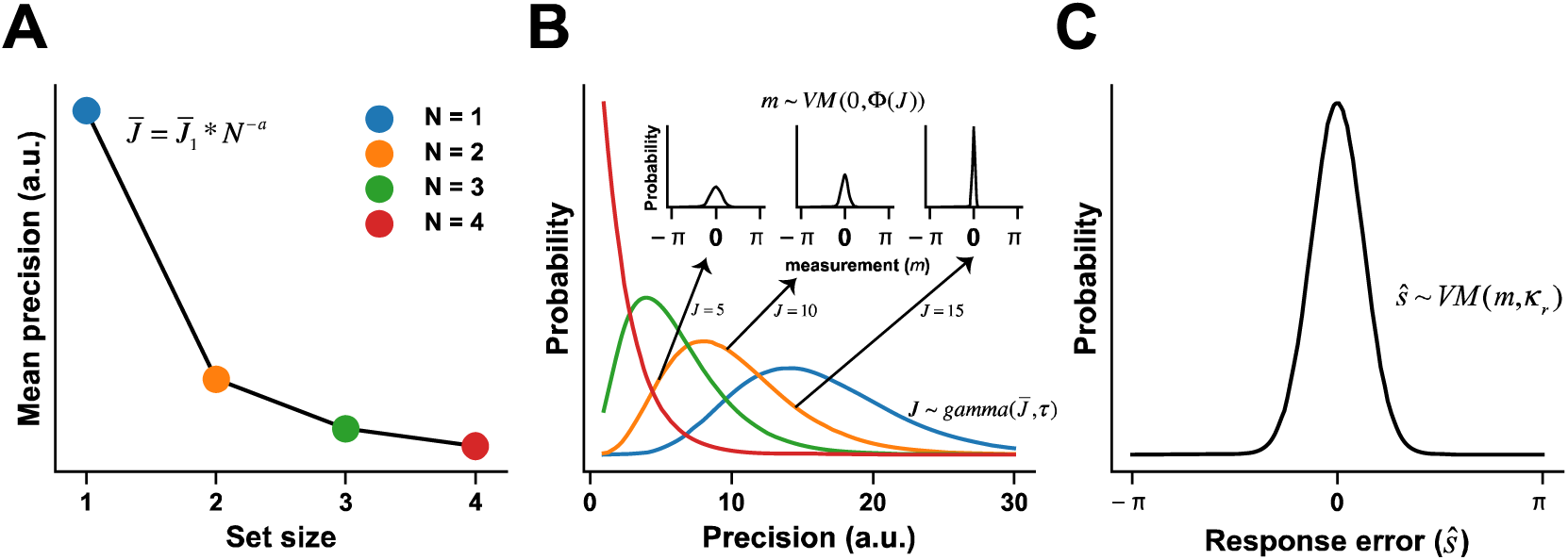
Variable-precision model of VWM. ***A***. Resource decay function. The VP model assumes that the mean resource 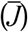 for processing a single item declines as a power function of set size *N*, a trend characterized by two free parameters—initial resources 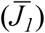 and decaying exponent (*a*). ***B***. The resources across items or trials follow a gamma distribution with the mean resource 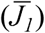 determined by the resource decay function (panel A) and the resource allocation variability (τ). Larger amounts of resources (*J*) indicate higher precision and therefore generate narrower von Mises distributions (three small axes indicating the precision equals to 5, 10 and 15 respectively) of stimulus measurement (*m*). The widths of the von Mises distributions indicate the degree of trial-by-trial sensory uncertainty. ***C***. The eventual behavioral choice given the internal stimulus measurement (*m*) is also uncertain, following a von Mises distribution with the choice variability (κ_*r*_) ^50^. In the VP model, initial resources 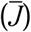, decaying exponent (*a*), resource allocation variability (τ) and choice variability (κ_*r*_) are four free parameters to estimate (see details in SI and van den Berg *et al*. ^48^). All numbers here are only for illustration purposes and not quantitatively related to the model fitting results in this paper.

We compared all seven models using the Akaike information criterion (AIC) and the Bayesian information criterion (BIC) ^46,47^ in each subject. In the color experiment, we found the VP model as the best-fitting model for over 84% of subjects in the HC group under both metrics (Fig. 3), replicating previous results in normal subjects ^48,49^. Most importantly, the VP model was also the best-fitting model for over 90% of subjects in the SZ group (Fig. 3). This result indicates that both groups use the same observer model to perform the task.

The results were replicated in the orientation delay-estimation task. We found that among 9 SZ subjects, the VP model was the best in 7 subjects using AIC and in all 9 subjects using BIC. Similarly, in 26 HC subjects, the VP model outperformed than other models in 21 and 24 subjects using AIC and BIC, respectively.

The superior performance of the VP model indicates the important role of variable precision in VWM processing. It is worth noting that the VP model, as the best-fitting model, does not include a capacity term. As such, we cannot conclude that the two groups must have the same capacity. But this result highlights the importance of systematic model comparisons before comparing group differences on model parameters.

### Larger resource allocation variability in SZ

Analyses above have established that HC and SZ employ the qualitatively same observer model to complete the VWM task. Their behavioral differences thus should arise from the differences on some parameters in the observer model. We next compared the fitted parameters of the VP model in the two groups. In the color experiment, we found that the two groups had comparable resource decay functions (Fig. 5A, initial resources, t(119) = 0.689, p = 0.492, d = 0.125; decaying exponent, t(119) = 1.065, p = 0.289, d = 0.194), indicating a similar trend of diminished memory resources as set size increases. SZ, however, had larger variability in allocating resources (Fig. 5B, resource allocation variability, t(119) = 4.03, p = 9.87 × 10^−5^, d = 0.733). Furthermore, the VP model explicitly distinguishes the variability in processing items and the variability in exerting a behavioral choice (e.g., motor or decision noise). We found no significant group difference in the choice variability (Fig. 5C, t(119) = 1.7034, p = 0.091, d = 0.31), excluding the possibility that the atypical performance of SZ arises from larger variability at the choice stage.

**Figure 5.**
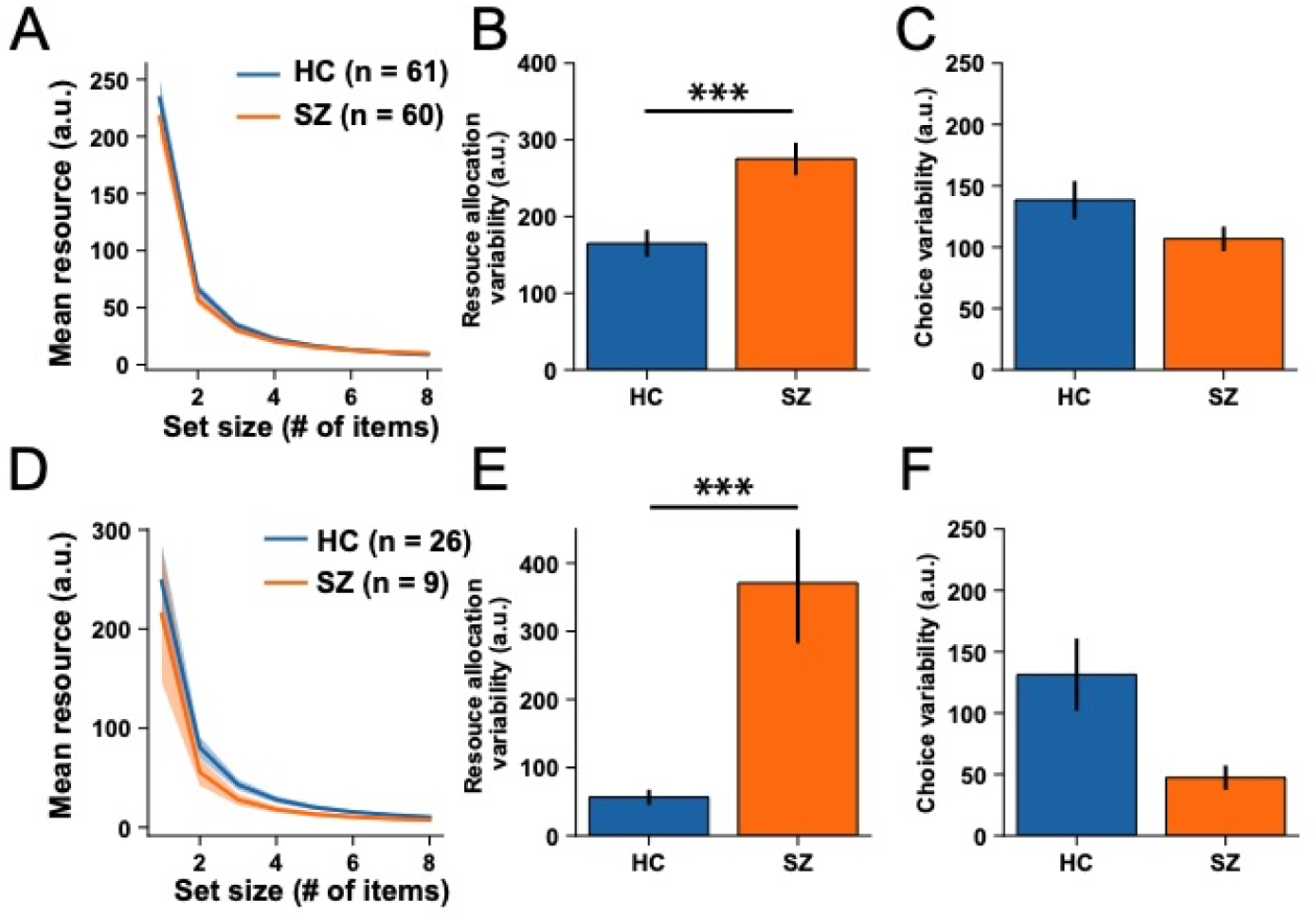
Fitted parameters of the VP model in color (***panels A-C***) and orientation (***panels D-F***) experiments. In the color experiment, no significant group differences are noted between two groups in resource decay functions (***panel A***), and choice variability (***panel C***). SZ have larger resource allocation variability than HC (***panel B***). The results are replicated in the orientation delay-estimation experiment (***panels D-F***). In panels A&D, the individual resource decay functions are computed by 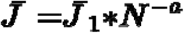, where *N* is the set size, and *a* are the estimated initial resources and the decaying exponent. The solid lines represent the averaged resource decay functions across subjects. The shaded areas in panels A and D, and all error bars in the other panels represent ±SEM across subjects. Significance symbol conventions are ***: p < 0.001.

Highly consistent results were again discovered in the orientation experiment: resource allocation variability was statistically higher in the SZ subjects (t(33) = 5.833, p = 1.576^-6^, d = 2.256), and no significant group differences were detected in other parameters (initial resources, t(33) = 0.437, p = 0.665, d = 0.169; decaying exponent, t(33) = 0.145, p = 0.886, d = 0.056; choice variability, t(33) = 1.651, p = 0.108, d = 0.638).

These results suggest that, although the two groups have on average the same amount of memory resources across different set size levels, SZ allocate the resources across items or trials in a more heterogeneous manner, with some items in some trials receiving considerably larger amounts and vice versa in other cases. This is unnatural since the probe was randomly chosen among all presented items with an equal probability. The more common strategy therefore should be to assign an equal amount of resources to every item and in every trial to tackle the unpredictable target. Note that in the condition of set size equal to 1 (i.e., only one item is presented), resource allocation cannot vary across items but may still vary across trials, leading to worse performance in SZ. We further quantitatively confirmed that increased resource allocation variability indeed leads to larger behavioral response errors (see Supplementary Fig. S4).

#### Resource allocation variability predicts the severity of schizophrenic symptoms

We next turned to investigate whether the results from the VP model can predict clinical symptoms. In both tasks, a set of correlational analyses was carried out to link the estimated resource allocation variability to the schizophrenia symptomatology in each subject. BPRS, SANS, and SAPS questionnaires were administered for each subject in the color experiment (Table 1 in Methods). The Positive and Negative Syndrome Scale (PANSS)^51^ was administered for each subject in the orientation experiment. PANSS includes positive symptomatology, negative symptomatology, as well as general psychopathology symptoms (Table 2 in Methods).

**Table 1.** Demographics and clinical information of people with schizophrenia (SZ) and healthy control subjects (HC)

|  | SZ (N = 60) |  | HC (N = 61) |  |
| --- | --- | --- | --- | --- |
|  | Mean | SD | Mean | SD |
| Age | 35.67 | 6.58 | 33.82 | 9.90 |
| range | 23-48 | n/a | 21-57 | n/a |
| Female/male | 31/29 | n/a | 29/32 | n/a |
| Inpatient/outpatient | 33/27 | n/a | n/a | n/a |
| Subject's education (in years) | 12.03 | 2.24 | 15.13 | 3.70 |
| Paternal education (in years) <sup>a</sup> | 9.89 | 2.53 | 9.76 | 2.95 |
| Maternal education (in years) | 9.62 | 2.91 | 9.29 | 3.63 |
| BPRS | 27.25 | 6.27 | n/a | n/a |
| SAPS | 5.77 | 7.02 | n/a | n/a |
| SANS | 24.43 | 11.45 | n/a | n/a |

<sup>a</sup> Average of mother's and father's years of education
BPRS: Brief Psychiatric Rating Scale<sup>80</sup>; SAPS: Scale for the Assessment of Positive Symptoms<sup>82</sup>; SANS: Scale for the Assessment of Negative Symptoms<sup>81</sup>.

**Table 2.** Demographics and clinical information of the subjects in the orientation delay-estimation task.

|  | SZ (N = 9) |  | HC (N = 26) |  |
| --- | --- | --- | --- | --- |
|  | Mean | SD | Mean | SD |
| Age | 39.33 | 9.35 | 21.31 | 1.94 |
| range | 27-56 | n/a | 19-26 | n/a |
| Female/male | 0/9 | n/a | 16/26 | n/a |
| Inpatient/outpatient | 9/9 | n/a | n/a | n/a |
| Subjects' education (years) | 9 | 2.12 | 14.12 | 1.63 |
| PANSS - positive | 11.33 | 4.33 | n/a | n/a |
| PANSS - negative | 22 | 8.14 | n/a | n/a |
| PANSS - general | 31.78 | 14.10 | n/a | n/a |

PANSS: The Positive and Negative Syndrome Scale<sup>51</sup>

In the color experiment, we noticed that the estimated resource allocation variability values of the SZ subjects correlated with their BPRS scores (Fig. 6A, r = 0.259, p = 0.045) and SANS scores (Fig. 6B, r = 0.302, p = 0.019). No significant correlation was noted on the SAPS scores (Fig. 6C, r = −0.121, p = 0.358). In the orientation experiment, we also found that resource allocation variability significantly correlated with the severity of general psychopathology symptoms (Fig. 6D, r = 0.717, p = 0.03) and negative symptoms (Fig. 6E, r = 0.882, p = 0.002), but not positive symptoms (Fig. 6F, r = 0.551, p = 0.124).

**Figure 6.**
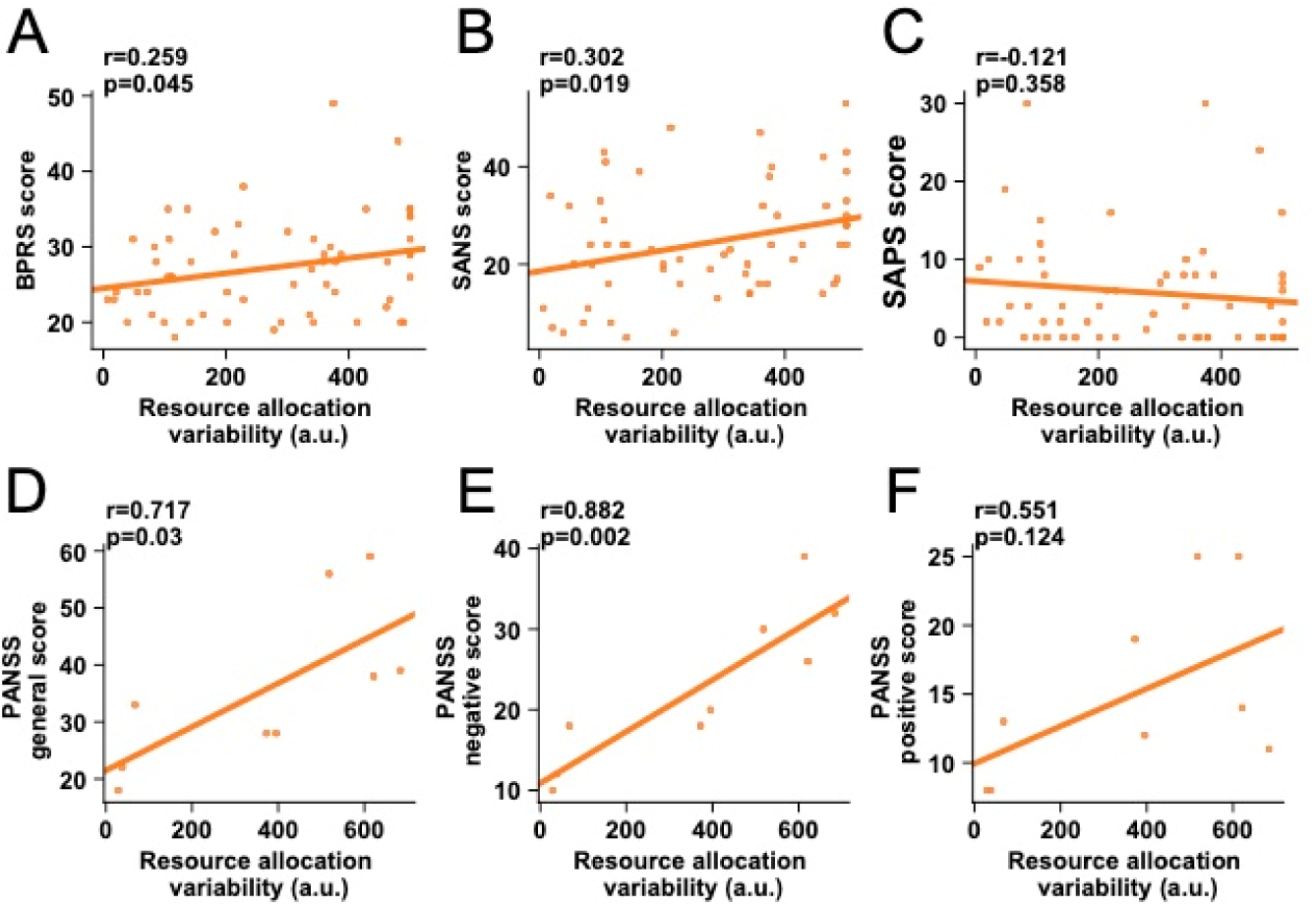
Individual differences in resource allocation variability predict the scores in symptom assessments. In the color experiment, estimated resource allocation variability values in the SZ group significantly correlates with their scores on BPRS (***panel A***) and SANS (negative symptoms, ***panel B***) but not on SAPS (positive symptoms, ***panel C***). In the orientation experiment, resource allocation variability significantly correlates with the PANSS general scores (***panel D***), PANSS negative scores (***panel E***). This correlation is not significant for PANSS positive scores (***panel F***).

Using different behavioral tasks and different measurements of schizophrenia symptomatology, we found consistent results that the resource allocation variability is linked to negative but not positive symptoms. These results suggest that resource allocation variability not only is the key factor explaining VWM behavior in SZ but also can quantitatively predict the severity of clinically measured symptoms, highlighting its promising functionality in future diagnosis and rehabilitations of schizophrenia.

#### Control experiments

To further exclude the potential confounding factors, we run two additional control experiments.

In the first control experiment, we tested the color perception of the subjects in a color estimation task. This task is similar to the color delay-estimation task except that there was no delay period and the probe was always shown on the screen such that subjects could directly see the probe when choosing colors on the color-wheel (see details in Supplementary Materials). We found that the individuals’ color perception thresholds cannot explain the group difference in resource allocation variability in Fig. 5B, indicating that the group differences are indeed memory related, not due to general worse perception in SZ.

In the second experiment, we aimed to exclude the possibility of imprecise model fitting due to low set size levels used in the color experiment. In order to recruit a large sample of subjects, we reduced the task difficulty and only tested two relatively low set size levels (i.e., 1/3) in the color experiment. One caveat of this approach is the possible imprecise model fitting because we did not challenge subjects’ ceiling performance. Although we obtained highly consistent results in the orientation experiment using more and higher set size levels, there still exist the possibilities that 1) the VP model might not be the best model, and more importantly, 2) the estimation of the model parameters (e.g., capacity parameter *K* in the VPcap model) might be inaccurate.

We further tested another large cohort of HC subjects (N=62) on the identical color delay-estimation task but used higher set size levels (i.e., 2, 4, and 6). Memorizing the colors of 6 objects is extremely challenging, and previous studies usually regarded 6 as the ceiling condition in a delay-estimation task ^37,52,53^. We performed the same analyses and found that the VP model was still the best-fitting model in over 70% of the subjects (Supplementary Fig. S3).

In sum, the three datasets (the color experiment, the orientation experiment, and the control color experiment) including a total of 149 HC and 69 SZ subjects demonstrate the converging result that the VP model is the best model for both HC and SZ subjects. This finding is unlikely due to some idiosyncratic experimental settings.

## Discussion

The mechanisms of VWM deficits in schizophrenia have been a matter of debate over the past few years. Decreased capacity has been widely proposed as the major cause of the deficits in SZ. In the present study, we re-examine this conclusion by comparing the performance of SZ and HC using several mainstream computational models of VWM proposed so far. We first establish that the VP model is the best model to characterize performance of both groups, indicating a qualitative similar internal process in both groups. We then further evaluate different components in the VP model and find that SZ have larger variability in allocating memory resources. These findings are highly consistent in two independent samples of subjects and in two independent behavioral tasks. Furthermore, individual differences in resource allocation variability predict variation of patients’ symptom severity, highlighting its clinical functionality. Taken together, our results propose for the first time that resource allocation variability is the key factor that limits VWM performance in schizophrenia.

A large body of literature has documented that SZ perform poorly in various forms of working memory tasks ^2,3,54,55^. The majority of the studies made the same conclusion: memory capacity is decreased in schizophrenia. However, through a careful examination of the literature, we find that the definition of capacity varies substantially across studies. Many studies directly equated worse performance with decreased capacity without quantitatively demonstrating how capacity modulates performance. For example, memory capacity was defined as the number of digits that can be recalled in the longest strand in digit span tasks ^12^. In N-back tasks, capacity was defined as the number of backs corresponding to a certain accuracy level ^14–16^. Moreover, the calculation of capacity resembled the d-prime metric in change detection tasks ^22–24,43,56^. The majority of these metrics are actually behavioral thresholds or accuracy related to capacity rather than direct quantifications of capacity. This is partly because we lack appropriate computational models for the majority of those tasks. The VP model is advantageous as it describes the generative process of the delay-estimation task and the change-detection task ^48^. As such, it allows to disassociate the effect of capacity from other VWM components. Note that here we only consider resource, capacity, resource allocation, and choice as the dimensions of modeling. Some studies suggest the existence of “binding error” in VWM ^42,57^. Van den Berg et al 2014 ^49^ explored a large space of model parameters and found that “binding error” only account for small fraction of VWM limitation. As such we did not consider “binding error” in this paper. Also, all computational models we used in this study are mostly specific to the delay-estimation task. It is possible that the VWM deficits in schizophrenia manifest differently in different tasks. Future studies are needed to test whether larger resource allocation variability can also account for VWM deficits of SZ in other WM tasks.

Only a few studies have quantitatively estimated capacity and precision in schizophrenia. Gold et al ^25^ employed the same color delay-estimation task to that in our study and estimated individual’s capacity and precision using the MIX model. Results in that study echoed the decreased-capacity theory. The MIX model, however, does not consider two important factors. First, the model assumes an equal precision across items in memory. Second, the model does not separate the variability for processing stimuli (i.e., sensory uncertainty, κ in Supplementary Eq. S5) and the variability in exertion of a choice (i.e., choice uncertainty, κ_*r*_ in Supplementary Eq. S6). This distinction is important since it highlights different types of uncertainty in encoding and decoding stages of VWM. Mathematically, these two types of uncertainty can be distinguished by manipulating set size since the encoding variability depends on set size but the choice variability does not. The issues of the MIX model have been symmetrically addressed in recent work ^58^.

Compared with capacity and precision—the two diagnostic features of VWM—resource allocation variability emerges as a new concept in VWM. It refers to the heterogeneity of allocating resources across multiple items and trials. We speculate that resource allocation variability might reflect the stability of attentional control when the brain processes multiple objects. First, it has been shown that attention and WM are two core components of executive control and tightly linked with each other ^59,60^. Second, schizophrenia is known to have deficits in top-down attentional modulation ^55,59^. Particularly, recent studies discovered the phenomenon of spatial hyperfocusing in schizophrenia patients^19,61–63^. If schizophrenia patients overly attend to one item and ignore others in the memory encoding stage, unbalanced resource allocation will likely occur. Note that SZ also exhibit worse performance when only one target is presented. In this condition, resources do not vary across items (i.e., only one time) but could vary across trials. Also, we want to emphasize that such variability is not equivalent to attentional lapse. A higher attentional lapse rate may lead to overall fewer resources, a phenomenon we did not observe in this study.

What are the neural mechanisms of this resource allocation variability? Recent neurophysiological studies proposed that the neural representation of a stimulus may follow a doubly stochastic process ^64,65^, which suggests that the variability in encoding precision is a consequence of trial-to-trial and item-to-item fluctuations in attentional gain ^32,48,66^. A recent study combined functional magnetic resonance imaging and the VP model, showing that the superior intraparietal sulcus (IPS) is the cortical locus that controls the resource allocation ^67^. Interestingly, schizophrenia patients have been known to have IPS deficits ^68^. Note that besides top-down factors, we cannot rule out the contribution of bottom-up neural noise in perceptual and cognitive processing ^64,65^, as found in several other mental diseases ^33–36^. Also, a recent study found that variable precision is even more likely to be driven by stimulus-specific effects ^69^.

The current results also reveal links between resource allocation variability and patients’ negative symptoms, but not positive symptoms (Fig. 6). These findings are consistent with several experimental and meta-analysis studies claiming dissociable mechanisms underlying the cluster of negative symptoms versus that of positive symptoms ^70–73^. More broadly, a growing collection of evidence suggests that visual perceptual deficits in schizophrenic patients are more likely to link to negative rather than positive symptom severity ^74–78^. Negative symptoms in turn might produce improvised social functioning. Humans depend heavily on VWM to interact with multiple agents and complete social tasks. Deficits in distributing processing resources over multiple agents therefore might cause disadvantages in social cognition.

In conclusion, our study proposes a new explanation that the resource allocation variability accounts for the atypical VWM performance in schizophrenia. This view differs from the decreased-capacity theory and provides a new direction for future studies that attempt to promote diagnosis and rehabilitation for schizophrenic patients.

## Methods

### Ethics Statement

All experimental protocols were approved by the institutional review board at the East China Normal University. All research was performed in accordance with relevant guidelines and regulations. Informed written consent was obtained from all participants.

### Color delay-estimation experiment

#### Subjects

61 HC and 60 SZ participated in the experiment. SZ were clinically (symptom and medication) stable inpatients (N = 33) and outpatients (N = 27) who met DSM-IV criteria ^79^ for schizophrenia. Patients having the history of any other mental or neurological disorders were excluded. All patients were receiving antipsychotic medication (2 first-generation, 43 second-generation, 15 both). Symptom severity was evaluated by the Brief Psychiatric Rating Scale (BPRS) ^80^, the Scale for the Assessment of Negative (SANS) and Positive Symptoms (SAPS) ^81,82^. HC were recruited by advertisement. All HC had no current diagnosis of axis 1 or 2 disorders as well as no family history of psychosis nor substance abuse or dependence. All subjects are right-handed with normal sight and color perception.

The two groups were matched in age (t(119) = 1.58, p = 0.118, d = 0.284), gender (31 females and 29 males) and education level of parents (t(119) = 0.257, p = 0.798, d = 0.047). Inevitably, the SZ had fewer years of education than the HC (t(119) = 5.51, p = 2.09 × 10^−7^, d = 1.00). The detailed demographic information is summarized in the Table 1.

#### Stimuli and Task

The subjects sat 50 cm away from an LCD monitor. All stimuli were generated by Matlab 8.1 and Psychtoolbox 3 ^83,84^, and then presented on a LCD monitor.

In the color delay-estimation task (Fig. 1A), each trial began with a fixation cross presented at center-of-gaze for a duration randomly chosen from a sequence of 300, 350, 400, 450 and 500 ms. Subjects shall keep their fixation on the cross throughout the whole experiment. A set of colored squares (set size = 1 or 3) was shown on an invisible circle with 4° radius. Our pilot experiment showed that SZ patients exhibit a high dropout rate if the task is longer than 30 mins or too hard (i.e., set size > 4). We therefore reduced the difficulty of the color task to set size level 1 and 3. The sample array lasted 500 ms. Each square was 1.5° × 1.5° of visual angle. Their colors were randomly selected from the 180 colors that are equally distributed along the wheel representing the CIE L*a*b color space. The color wheel was centered at (L = 70, a = 20, b = 38) with a radius of 60 in the color space ^37^. The sample array then disappeared and was followed by a 900 ms blank period for memory retention. After the delay, an equal number of outlined squares were shown at the same location of each sample array item, with one of them bolded as the probe. In the meantime, a randomly rotated color wheel was shown. The color wheel was 2.1° thick and centered on the monitor with the inner and the outer radius as 7.8° and 9.8° respectively. Subjects were asked to choose the remembered color of the probe by clicking a color on the color wheel using a computer mouse. Subjects shall choose the color as precisely as possible and response time was not constrained. Every subject completed 2 blocks for the set size 1 and 3, respectively. The order of the two blocks was counterbalanced across subjects. Each block had 80 trials. The difference between the reported color and the true color of the target is considered as the response error.

### Orientation delay-estimation experiment

#### Subjects

Data from 26 HC and 9 SZ were obtained in the orientation delay-estimation task (see Table 2 for subject information). The data from the HC subjects has been previously presented in ref. ^41^. SZ were all clinically stable inpatients who met the DSM-IV criteria for schizophrenia^79^. Patients having a history of any other mental or neurological disorders were excluded. All nine patients were receiving second-generation antipsychotic medication. The Positive and Negative Syndrome Scale (PANSS)^51^ was used to evaluate the psychotic symptoms of the patients. This scale includes positive symptomatology, negative symptomatology, as well as general psychopathology symptoms. HC subjects were recruited by advertisement. All HC had no current diagnosis of axis 1 or 2 disorders as well as no family history of psychosis nor substance abuse or dependence. All subjects are right-handed with normal sight.

#### Stimuli and Task

Stimuli were presented on a 60 Hz LCD monitor through Matlab psychophysics Toolbox (Version 3). In the sample array, all items were shown on an invisible circle with 10° radius. The length and width of each item in the sample array were 3.7° and 0.6° respectively. Then orientations of the bars were randomly chosen from 1° to 180°. The procedure of this task was similar to the color delay-estimation task, except that the sample array was presented for 200 ms (Fig. 1B). In the probe array, one of the sample bars appeared as the probe, and subjects were required to adjust the orientation of the probed bar using a computer mouse.

The HC subjects completed 5 blocks with 100 trials in each block. The set size of each trial was randomly chosen from 1, 2, 3, 4, 6. The SZ patients complete the experiment in three visit sessions. In each visit, they were asked to finish 10 blocks with 40 trials in each block. The set sizes were 1, 2, 4, 6 and counter-balanced across blocks. We analyzed the behavioral performance using only the data of set size levels 1, 2, 4, 6 because they were the conditions shared by both groups. For model fitting, we included all set size levels (1/2/3/4/6) of the HC group and compared the likelihood, AIC, and BIC per trial to compensate the trial difference between groups and subjects. One patient only finished 9 blocks in the second visit and another patient only finished 8 blocks in the first visit. In total, seven SZ subjects completed 300 trials for each set size, one SZ completed 290 trials and one SZ completed 280 trials.

## Data availability statement

The data are available from the corresponding author upon reasonable request.

## Acknowledgments

We thank Zheng Ma, Ting Qian and Haojiang Ying for their invaluable comments on the manuscript. This work was supported by the National Social Science Foundation of China (17ZDA323), the Shanghai Committee of Science and Technology (19ZR1416700, 17JC1404101, 17JC1404105) to Y.K.

## Author contributions

Y. K., Y. Z. and X. R. designed the experiments. X. R. and L. Z. performed the experiments. Y. Z., T.M. and R-Y. Z. analyzed the data, Y. Z and R-Y. Z. wrote the manuscript in consultation with Y. K. and L. Z.

