## Supplemental Materials for "Atypically larger variability of resource allocation accounts for visual working memory deficits in schizophrenia"

**This PDF file includes:**

Computational models of VWM

Intuitive model explanations

Model fitting and comparisons

Control color delay-estimation experiment and statistical results

Control color estimation experiment and statistical results

Supplementary Figures. 1 to 4

References

**COMPUTATIONAL MODELS OF VISUAL WORKING MEMORY**

**Variable-precision model**. The variable-precision (VP) model has been shown as the state-of-the-art computational model of VWM. Details of the VP model have been documented in several previous studies ^1,2^ and the model codes are publicly available (<http://www.cns.nyu.edu/malab/resources.html>).

The VP model assumes a resource decaying function describing the decreasing trend of mean memory resource (
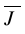
) assigned to individual items as the set size (*N*) increases ^3,4^:

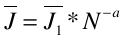
 , (S1)

where
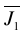
 is the initial resources when only 1 item (*N* = 1) should be memorized and *a* is the decaying exponent. The key component of the VP model is that the memory resources
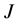
 across items and trials follow a Gamma distribution with the mean
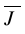
and the scale parameter
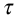
:

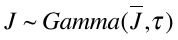
 , (S2)

Intuitively, a larger
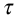
 indicates a more uneven distribution of memory resources across items or trials, with some items in some trials receiving a larger amount of resources while others receive comparative fewer. Note that a larger amount of memory resource produces a higher precision. Thus, we do not explicitly distinguish resource and precision and denote them as *J.* Defining precision as Fisher information ^5^, precision
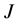
 can be linked to the variance of the von Mises distribution of sensory measurement:

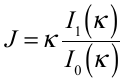
, (S3)

where
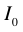
 and
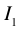
 are modified Bessel functions of the first kind of order 0 and 1 respectively, with the concentration parameter
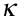
. Eq. S3 specifies a one-on-one mapping between precision
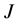
 and variance
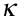
. We can rewrite their relationship as:

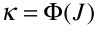
 , (S4)

where
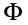
 is the mapping function. The distribution of sensory measurement (*m*) given the input stimulus (*s*) can be written as:

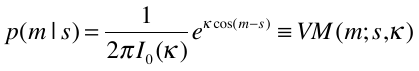
, (S5)

We further assume that the reported color (
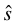
) by participants also follows a von Mises distribution:

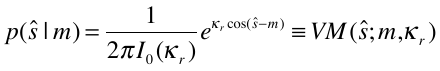
, (S6)

where
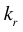
 represents the variability at the choice stage.

Given the four free parameters and stimulus color
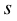
 in a trial, we can derive the probability of the observed response in a trial by marginalizing over sensory measurement
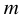
 and variable precision
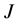
:

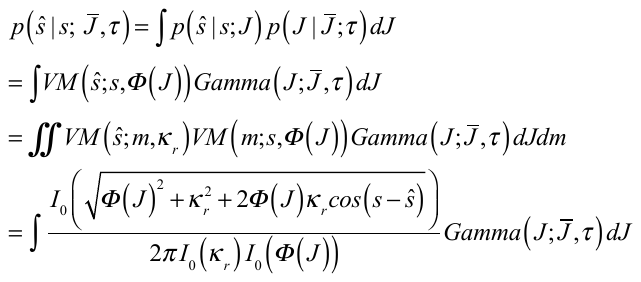
 ，(S7)

Note that in Eq. S7, sensory measurement (*m*) can be analytically eliminated. Since precision
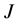
is a random variable across items and trials, we sampled it 10000 times from the Gamma distribution with mean
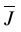
 and scale parameter
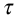
. Note that van den Berg *et al*. ^1^ confirmed that 500 samples are enough in the model fitting. We then used all the samples to calculate response probability in each trial.

Taken together, this VP model has four free parameters:
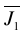
, *a*,
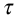
and
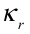
.

**Variable-precision-with-capacity model**. The variable-precision-with-capacity (VPcap) model inherits all parameters and the structure of the VP model above, except that an additional capacity parameter (*K*) is introduced to estimate the memory capacity of individuals. If the set size *N* is smaller than capacity *K*, the VPcap model is identical to the VP model. If the set size *N* exceeds the capacity *K*, the model assumes that the probe is stored in the VWM with the probability *K*/*N*, and out of memory with the probability 1- *K*/*N*. In the latter case, a participant randomly guesses a color. The response probability therefore can be written as:

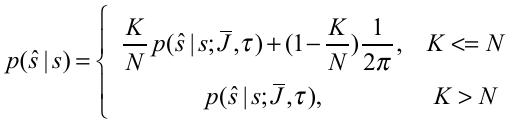
, (S8)

where
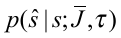
 is defined in Eq. S7. In essence, the VPcap model is a mixture model of the VP model and a random guessing process when the set size exceeds the participant’s capacity. The VPcap model has five parameters, four as the same in the VP model and the additional capacity parameter (*K*).

**Item-limit model***.* The item-limit (IL) model assumes no uncertainty in the sensory encoding stage such that the internal sensory measurement *m* is equal to the input stimulus *s*. But there exists choice variability from measurement *m* to the reported color (
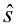
). Such choice variability does not vary across set size levels. The IL model also assumes a fixed capacity *K.* The response probability is:

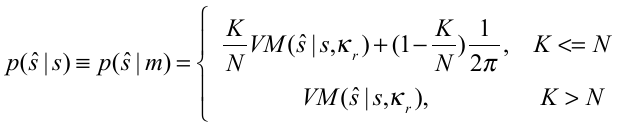
 , (S9)

The IL model has two free parameters: choice variability
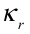
, and capacity *K*.

**Mixture model***.* The mixture model (MIX) has been used in previous clinical research ^6^. Similar to the IL model, the MIX model only assumes the uncertainty from stimulus *s* to the reported color (
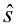
) and a fixed capacity *K.* The difference is that the uncertainty (
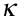
) reflects both sensory noise and choice variability, and thus the uncertainty is set-size dependent (each set size has one
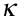
). The response probability can be written as:

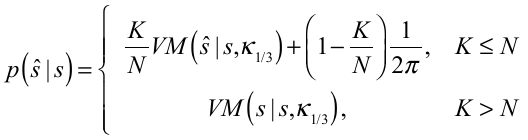
, (S10)

where and denote the uncertainty for set size 1 and 3, respectively. The MIX model has three parameters: uncertainty levels
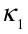
 and
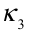
, and capacity *K.*

**Slots-plus-averaging model**. The slots-plus-averaging (SA) model was originally proposed in ^7^ and further elaborated in ^1^. Unlike the IL model, the SA model acknowledges the presence of noise in the sensory encoding stage. However, the memory resources are discrete chunks, and a single chunk or multiple chunks can be assigned to one item. For one item, the SA model assumes Eq. S4 still holds as the relationship between the resource assigned to that item and the width of the von Mises distribution:

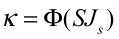
 , (S11)

where *S* is the number of chunks and *J_s_* is the resource of one chunk. The SA model also assumes a capacity *K*.

When *N* > *K*, an item should receive either 0 or 1 chunk. Then the allocation should be similar to the IL model. the response distribution should be a mixture of a uniform and a von Mises distributions:

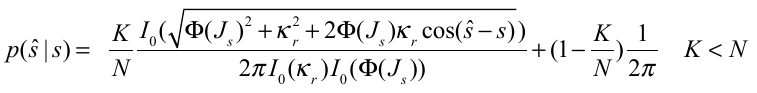
, (S12)

When N ≤ K, some items receive either one or more chunks. Assuming that the resource chunks should be assigned as equally as possible across items, the *S* can be calculated as:

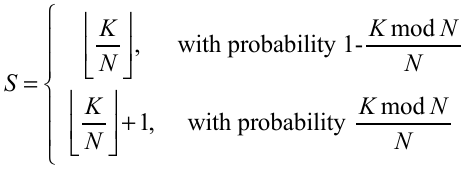
, (S13)

where
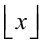
 represents the *floor* function in Matlab. The corresponding concentration parameter of von Mises distributions can be computed by Eqs. S11&13:

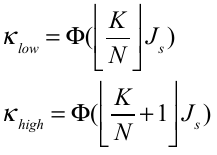
 (S14)

The response probability in the SA model can be written as:

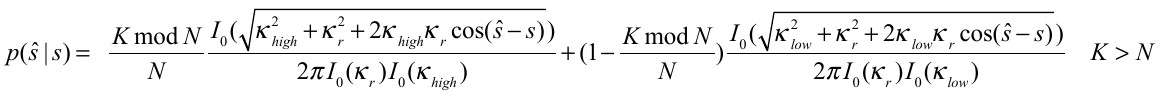
, (S15)

The SA model has three free parameters: unit resource *J_s_*, choice variability
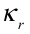
, and capacity *K.*

**Cosine slots-plus-averaging model**. A recent paper ^8^ suggests that a modified version of the SA model, dubbed cosine slots-plus-average model (cosSA), outperformed the VP model to explain the delay-matching VWM behavior. To enhance the generality of our study, we also followed that work and included this model. Briefly, the cosSA model assumes that the unit memory precision is stimulus-dependent and exhibits a cosine-like periodic fluctuation:

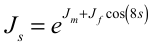
, (S16)

where
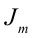
and
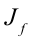
 describe the fluctuation of unit memory precision (
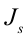
) as a function of stimulus *s*. Note that the frequency of the cosine function was derived from the cosine-shaped bias found in our empirical data. We can convert precision
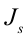
 to the width of von Mises distributions
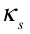
according to Eq. S4. According to capacity *K*, the discrete memory resource allocation is described as Eq. S11-S14. Moreover, the cosSA model also assumes the response bias is periodic:

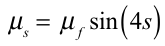
, (S17)

where
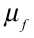
 adjusts the magnitude of the bias. The probability of a response given the stimulus can be described as:

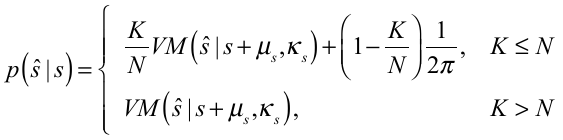
 , (S18)

The cosSA model has four free parameters:
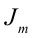
,
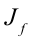
,
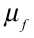
 and capacity *K*.

**Equal-precision model**. The equal-precision (EP) model is very similar to the VP model, except that an equal amount of resources is assigned to every item and in any trial. Namely, the Eq. S2 does not apply to the EP model. In the EP model, the resource assigned to one item declines as a power function (as Eq. S1). Then the resource at each set size level can be converted to the width of the von Mises distribution using (Eq. S4). The response probability is given by:

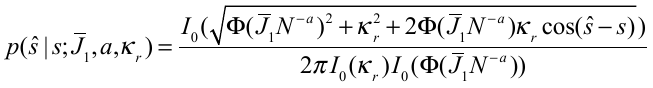
, (S19)

where *J_1_* is the resource when set size is 1 (initial resources). The EP model has three free parameters: initial resources
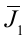
, decaying exponent *a*, and choice variability
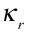
.

**INTUITIVE MODEL EXPLANATIONS**

Despite the mathematical details provided above, we further provide intuitive explanations for each model and highlight their differences based on cartoon illustrations in Fig. S1. Note that all stimuli are 0 because we transformed the reported color to recall errors in each trial.

**Item-limit model.** In the IL model (Fig. S1A), if the capacity *K* is larger than the set size N (e.g., N=2, K=3, the left panel), all items can enter working memory. The reported color follows a von Mises distribution with the mean as the color of the probed stimulus. If the capacity *K* is smaller than the set size *N* (e.g., N=2, K=3, the right panel), a probed stimulus can be stored within memory with probability *K*/*N* and out of memory with probability (1-*K*/*N*). If the probed stimulus is in memory, the same rule of von Mises distribution applies. If the probed stimulus is out of memory, a subject guesses a color (i.e., with probability 1/2π, the uniform distribution of guessing).

**Mixture model.** The mixture model (Fig. S1B) shares all components with the IL model. The key difference is that the IL model assumes the same von Mises distribution for both set size levels (i.e., the same width of the blue and the orange distributions in Fig. S1A), while the mixture model uses two von Mises distributions with different widths for the two set size levels (i.e., different widths of the blue and the orange distributions in Fig. S1B), to compensate the potential different level uncertainty associated with two set size levels. Thus, the mixture model has one additional free parameter than the IL model.

**Slot-plus-averaging and cosine slot-plus-averaging model.** The SA model regards memory resources as several discrete chunks (Fig. S1C). In the example of Fig. 1C, the subject has three (*K* = 3) chunks of resources and the blue cups stand for individual stimulus. If two stimuli are presented (i.e., two cups, set size = 2), the scenario in which the number of resource chunks is larger than the set size, two resource chunks are assigned to one cup and another chunk to the other cup. If the number of resources is smaller than the set size (e.g., four stimuli/cups), one cup will receive no resource, and the subject has to guess if this stimulus/cup is probed. The key difference between the SA model and the three models below is that the SA model assumes discrete resource chunks.

The cosSA model is a modified version of the SA model with three major changes ^8^. First, the unit memory precision is stimulus-dependent and follows a periodic function (see Eq. S16 and Fig. S1D). Second, it also includes a response bias that is also stimulus-dependent and periodic (see Eq. S17 above and Fig. S1D). Third, for simplicity, it does not include the response variability and only includes one uncertainty (i.g., encoding precision) in the processing.

**Equal-precision, variable-precision and variable-precision-with-capacity models.** The EP, VP, and VPcap models share one core assumption: memory resources are continuous, analogous to the amount of juice in a big mug (Fig. S1E). A subject assigns the juice (i.e., resources) into different cups (i.e., stimuli). In Fig. S1E, the orange cups stand for the mean juice amount an individual cup receives in each set size condition. We can imagine that, given the total amount of juice is fixed, the more cups (i.e., larger set size) the less juice on average each cup will receive. This is reflected by the diminishing average amount of juice as set size increases (also see Eq. S1).

Besides the core assumption of continuous resources, the three models have slightly different specifications (Fig. S1F). In Fig. S1F, all orange cups stand for the mean juice amount in each set size condition, and the blue cups stand for individual stimulus. The EP model assumes that in each set size condition, each cup receives an identical amount of juice (upper row in Fig. S1F). In the VP model, however, each cup receives a variable amount of juice even though their average amount is the same as in the EP model. Using two cups as an example, the average amount of juice might be 10 ml but one cup might have 9 ml and the other one has 11 ml. Whether the amount of juice in each cup varies is the key difference between the EP and the VP models. Moreover, both EP and VP models do not constrain the total number of cups. Therefore, a cup will more or less receive a little bit juice even though there is a large number of cups (middle row). In other words, both the EP and the VP models have no concept of capacity. In contrast, the VPcap model not only inherits the assumption of variable precision and but also constraints the maximal number of cups (i.e., capacity *K*) that can receive juice. If the total number of cups (i.e., *N* stimuli) is larger than the capacity *K*, some cups will receive no juice, and the subject has to guess the color of these stimuli.

**MODEL FITTING AND COMPARISONS**

The BADS optimization toolbox in MATLAB ^9^ was used to search the best-fit parameters that maximize the likelihood of response data in all trials. BADS has been shown to outperform other default nonlinear optimization algorithms in MATLAB, especially in the problems where gradients on loss function are not available or hard to compute ^9^. We fit all models separately in each participant. To avoid local minima, we repeated the optimization process with 20 different initial seeds that are equally spaced within a lower and an upper bound. Parameters bounds were set to be very broad to avoid bias. The parameters with the maximum likelihood value were used as the best-fit parameters for one subject.

We compared the performance of all models fitted in this study. Model comparisons were performed for both groups using both Akaike information criterion (AIC) and Bayesian information criterion (BIC) ^10,11^ metrics..

**CONTROL COLOR DELAY-ESTIMATION TASK**

We repeated the identical color delay-estimation task on 62 HC subjects (30 females, 19-24 years old). The stimuli and procedure were identical to the task described in the main text except that: 1) the sample array was presented for 200 ms; 2) the set sizes were 2, 4, and 6; 3) there were 100 trials in each set size.

We fitted all seven models to each individual’s data and performed the model comparison. We found that in the total 62 HC subjects, the VP model was the best in 45 and 54 subjects using the AIC and BIC metrics, respectively (see below Fig. S3).

**CONTROL COLOR ESTIMATION TASK AND RESULTS**

**Color estimation task.** Before the main VWM task, all subjects completed a task to measure their color perception ability. The task is identical to the VWM task except for two modifications. First, only one colored object was shown in the sample array. Second, in the probe array, the colored object appeared again on the screen. A subject needed to choose its color on the color wheel while looking at it. There was 1 block with 50 trials in this task.

**Color perception results between HC and SZ.** We used the circular standard deviation (CSD) of response errors (the circular distance between the original color and chosen color in a trial) to evaluate the performance in the color task. A significant group difference was found (t(119) = -2.095, p = 0.038, d = -0.38), suggesting in general worse color perception in SZ. But this result might also be explained by potential differences in choice variability (e.g., motor control). To exclude the potential confounding of color perception, we further set CSD from the color perception as a co-variate and repeat all statistical analyses (see below).

**VWM performance.** We added the CSD in the color perception task as a co-variate to VWM performance comparison of two groups. The repeated-measure ANCOVA (see the main text for details of variables) results again showed a worse VWM performance at higher set size level (F(1,119) = 100.676, p < 0.001, partial η^2^ = 0.46). The group was also significant (F(1,119) = 8.902, p = 0.003, partial $\eta^{2}$ = 0.070), indicating that HC’s performance was better than SZ’s. The interaction between set size and group was not significant (F(1,119) = 0.324, p = 0.570, partial $\eta^{2}$ = 0.003). Also, the color perception ability had no influence on VWM performance (F(1,119) = 0.285, p = 0.595, partial $\eta^{2}$ = 0.002). These results replicated the results from the main text.

**Fitted parameters of the VP model.** Univariate general linear models were used for comparing fitted parameters between the two groups. We regressed out the factor of color perception by setting. Same as results in the main text (Fig. 5), comparable resource decay functions (Fig. 5A, initial resources, F(1,119) = 0.376, p = 0.541, partial $\eta^{2}$ = 0.003; decaying exponent, F(1,119) = 0.573, p = 0.451, partial $\eta^{2}$ = 0.005) and choice variability (Fig. 5C, F(1,119) = 1.702, p = 0.195, partial $\eta^{2}$ = 0.014) between SZ and HC were found in this analysis. And SZ showed larger variability in allocating resources (resource allocation variability, F(1,119) = 15.112, p < 0.001, partial $\eta^{2}$ = 0.114).

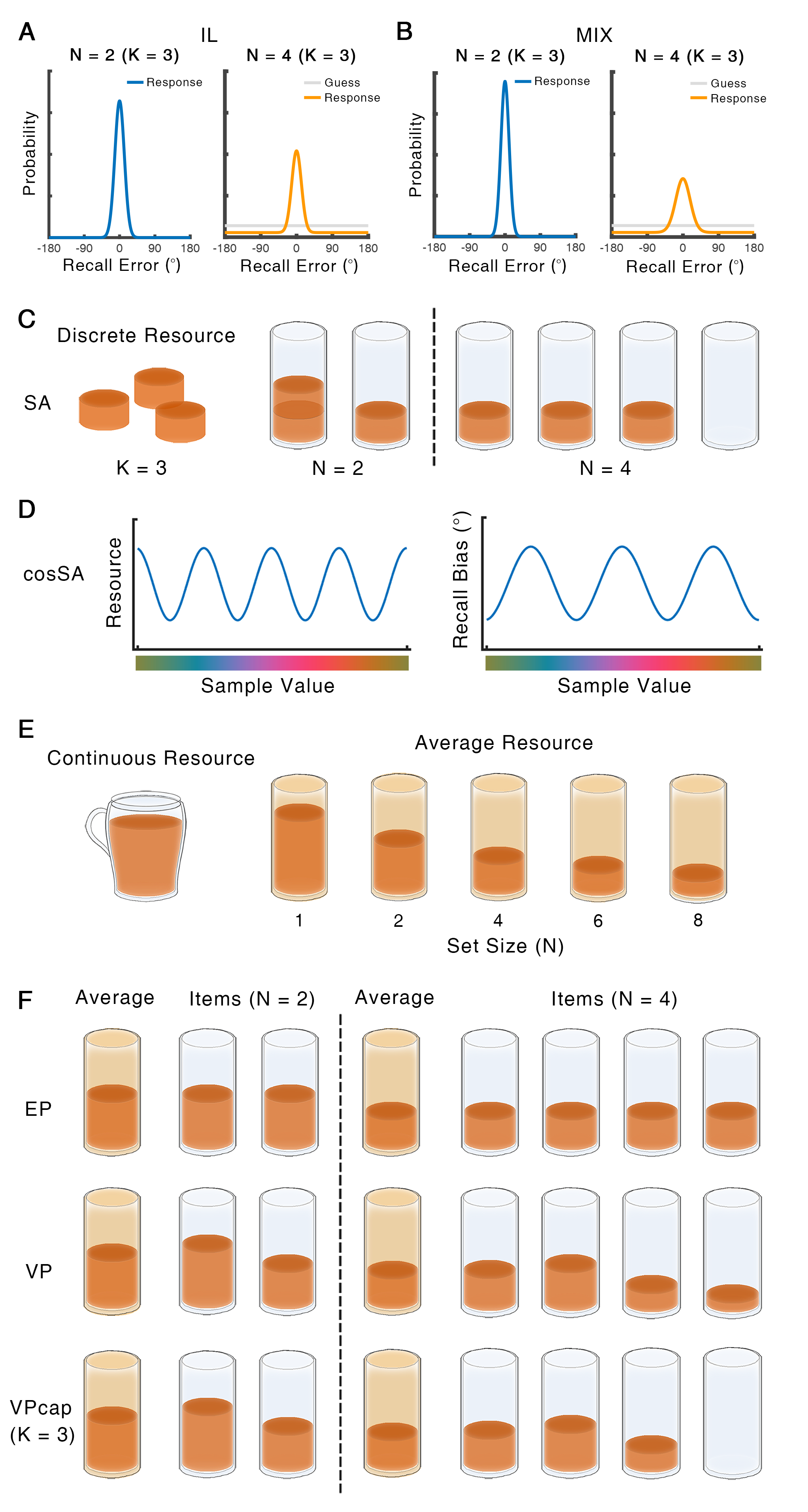

Supplementary Figure 1. Cartoon illustration of all computational models considered in this study. This figure aims to aid an intuitive understanding of the models. Detailed model explanations are in the section of ***intuitive model explanations***. ***A***. item-limit model; ***B***. MIX model; ***C***. the principle of discrete slots and the SA model; ***D***. cosSA model; ***E***. the principle of continuous resources; ***F***, EP, VP, and VPcap models. See the section of ***intuitive model explanations*** for detailed explanations.

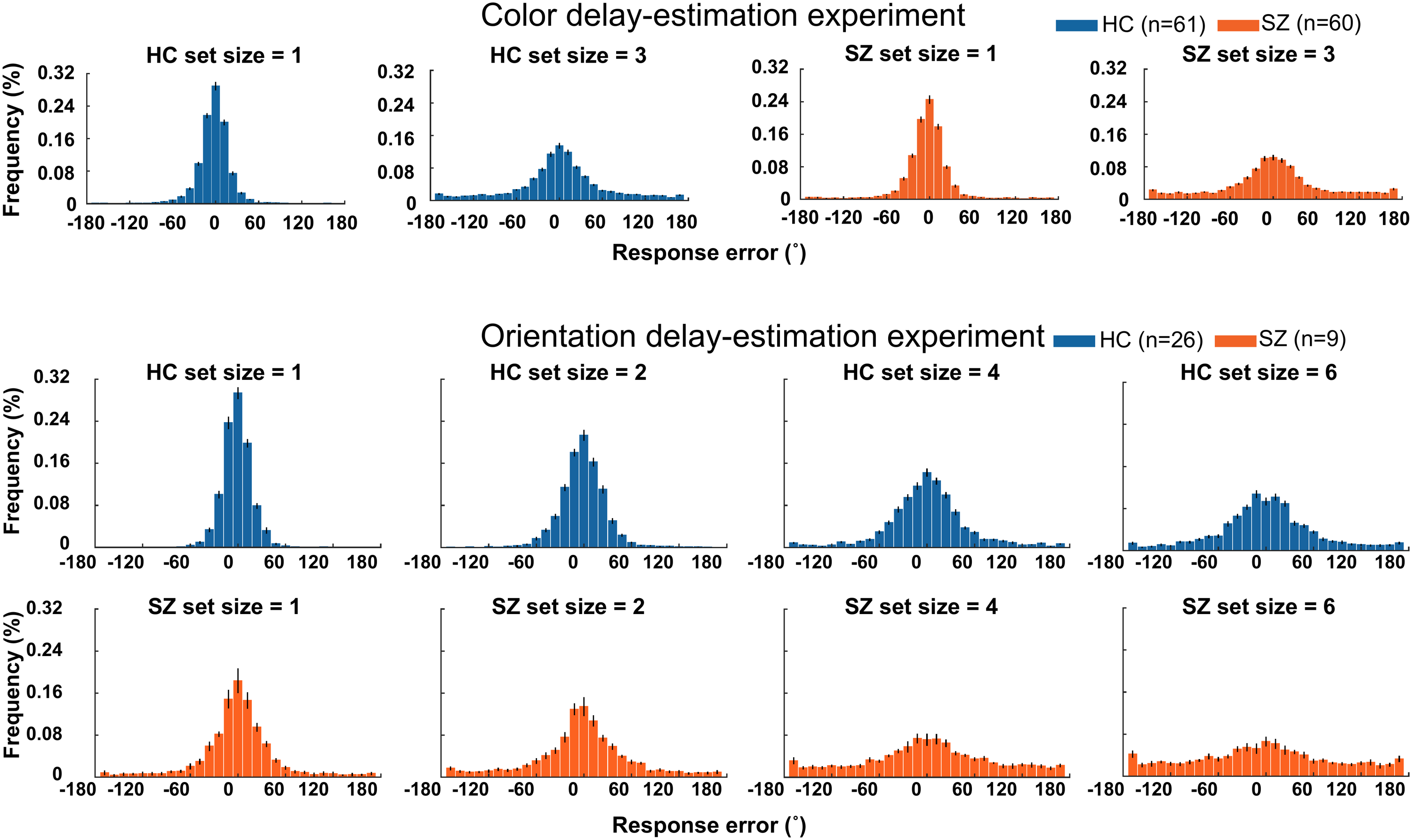

Supplementary Figure 2. Distributions of response errors across different set size levels in the two tasks. In all set size levels and in both tasks, the distributions of errors are wider in SZ subjects than that in HC subjects. Note that the standard deviations of these distributions are plotted in Fig. 2.

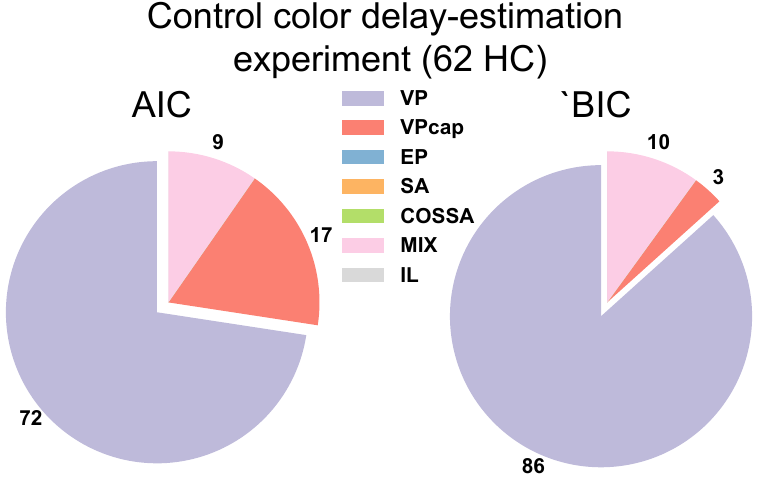

Supplementary Figure 3. Model comparisons on the data of 62 HC subjects tested in the control color delay-estimation task. Note that different from the color task in the main text, these subjects were tested on three set size levels (2/4/6). Model comparison results are consistent with those in the main text: the VP model is the best one when set size levels are increased.

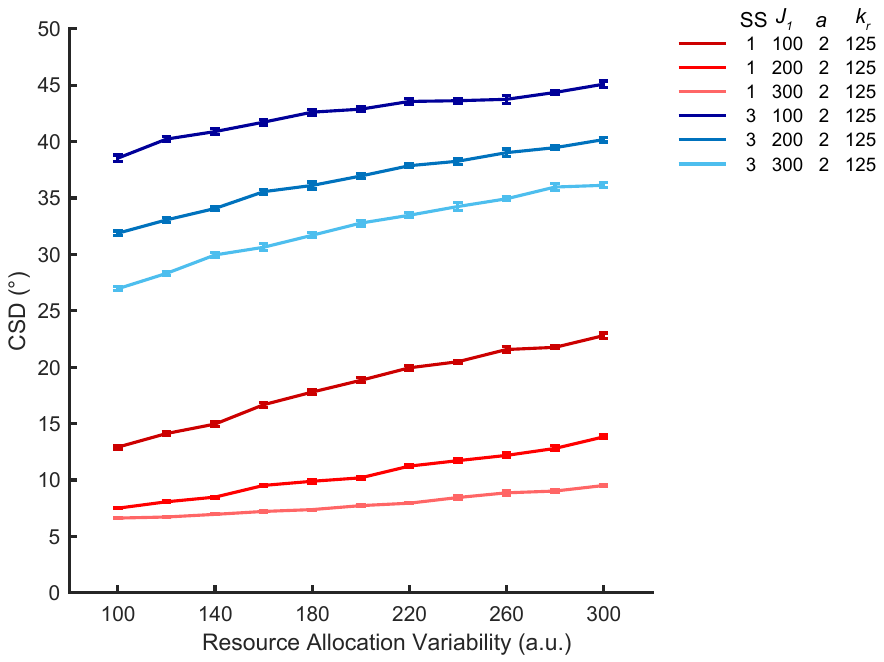

Supplementary Figure 4. Simulation of the behavioral consequences of increased resource allocation variability. Based on the VP model, we simulate 4000 behavioral responses in each parameter combination (SS: set size, $J_{1}$: initial resources, $a$: decaying exponent, $k_{r}$: choice variability). We systematically manipulate resource allocation variability and initial resource, and fix decaying exponent and choice variability to the group average of the fitted parameters of the HC group. Increased resource allocation variability leads to large response errors, indicating that low resource allocation variability is more optimal in this task.
